# BACTERIAL DIVERSITY IN ESTUARINE SEDIMENTS IN BRAZILIAN COASTAL: A FOCUS IN BACTERIAL RESISTANCE

**DOI:** 10.1101/2023.11.21.568096

**Authors:** Itana Almeida dos Santos, Carolina Oliveira de Santana, André Freire Cruz, Eddy José Francisco de Oliveira

## Abstract

Estuaries are final depositional environments of sediments transported along rivers. Therefore, they give a generalized dimension about the human effluents released in water systems. The development of molecular techniques allowed the study of environmental DNA (eDNA), which can provide a real dimension about the functioning of the microbial communities of those environments and the impacts of human activities. In this context, antimicrobial resistance is an emerging problem, caused by irresponsible use of antimicrobial drugs in human and animal health, which has contributed to a selection of bacteria resistant to these drugs. The objective of this study is to identify the main phyla of bacteria and to detect the presence of Antibiotic Resistance Genes (ARGs) in different mangroves environments. We observed significant differences between ARGs quantity in each point. Samples collected on Guaibim-2 and Una had bigger diversity of ARGs, while Guaibim-1 had less AGRs diversity. The predominant phyla in all of the samples was Proteobacteria, Actinobacteria and Firmecutes. We detected *Enterococcus faecium, Acinetobacter sp, Klebsiella pneumoniae, Enterococcus faecium* and *Enterobacter cloacae,* listed bacteria for priority surveillance for World Health Organization on points Una and Guaibim-2.

## INTRODUCTION

Mangroves are vegetal formations tolerant to brackish or salt water. They are situated along the coast of tropical and subtropical regions on transitional fields between river and sea, the estuaries[1,2]. These ecosystems offer many environmental services, such as fishing, coastal protection, wood production, indication of environmental risk, ecotourism, carbon sequester, bioremediation, protection against saline introduction into the groundwater, among others[1]. However, anthropic pressures in these environments (pollution, wood extraction, conversion of lands to unsustainable agriculture and aquaculture[3] and climate changes[4] are persistent and affect the state of conservation of mangroves along the world[2].

Formed with deposition of sediments that are carried along the river, soils of estuarine regions have high content of nutrients, therefore they are highly productive environments. These characteristics make this type of coastal formation hotspots of microbial diversity[5,6], wich has high biotechnological potential [5] The microbiome, the assembly of microorganisms of an environment, is responsible for important functions like organic matter decomposition, nutrient cycling and soil formation [7]. In this regard, microbial diversity of soils is influenced by biogeographic, human and relations between microorganisms[5].

Analysis of microbial diversity can be important bioindicators for environmental contamination[8],[9],[10][11]. Many studies has related larger heterogeneity of microbiome, or larger diversity of functional groups, to superior environmental quality[12][11], since this can make an ecosystem more dynamic and resilient, on hypothesis of occurrence of atypical events[13]. .

The study of the environmental DNA (eDNA) can elucidate many structural and functional aspects about microbial communities of a ecosystem. Through this perspective, it is possible to detect genetic material of microorganisms directly from environmental samples, without the need of cultivate them, wich can offer a real amplitude of microbial diversity.

Additionally, detection of antibiotic resistance genes (ARGs) through molecular analysis of eDNA revels anthropic impacts on bacterial diversity in soil[14],[15]. Indeed, ARGs can naturally occur on an environment, however, effluents of industrial, domestic or agropecuary systems can be a pression to the selection of these genes[14]. This occur because many of these effluents has antibiotic waste that can be a selective pressure for environmental bacterias and/or ARGs, which can be incorporated by bacterias on an environment through lateral transference of genes as transformation, transduction or through Mobile Genetic Elements, as transposons, integrons or plasmides[16].

Molecular techniques are effective for the study of many pathogens. Their dissemination for covid-19 pandemic was indispensable for tracking the origin of Sars-Cov-2, to understand its virulence and formulate plans to combating the pandemic [17]. Therefore, techniques like whole genome sequence, metagenomics and qPCR provide important data that are applied to comprehend resistance mechanisms of pathogens[18]. Environmental surveillance through eDNA can be an efficient strategy for detection of ARGs, heavy metals and other toxic substances. The integration of these techniques to public politics can subsidize strategies to avoid or mitigate the impact on environmental and human health[19],[20],[21],[22].

Water is one of the main via to transference of resistance mechanisms[23]. Therefore, ARGs can be identified both in aquatic systems and their sediments, but DNA abundance is larger in sediments[24].

Overall, starting from the problematic “Are there differences on microbial communities composition in estuarine sediments of different levels of anthropic impacts?”, our objectives are to investigate the microbial diversity in different environmentals and to detect ARGs and pathogenic bacterias associated to them through qPCR and new generation sequencing (NGS).

## METHODS

### Legal and ethical procediments

Legal or ethical permissions were not required for this study, since the samples were collected on public locations. The project is registered on SISGEN - National System of Genetic Patrimony Management, # A17C406

### Sample field

Valença city is located on state of Bahia (Figure 1), on the coastal zone of Atlantic Forest.

**Figure 1.**
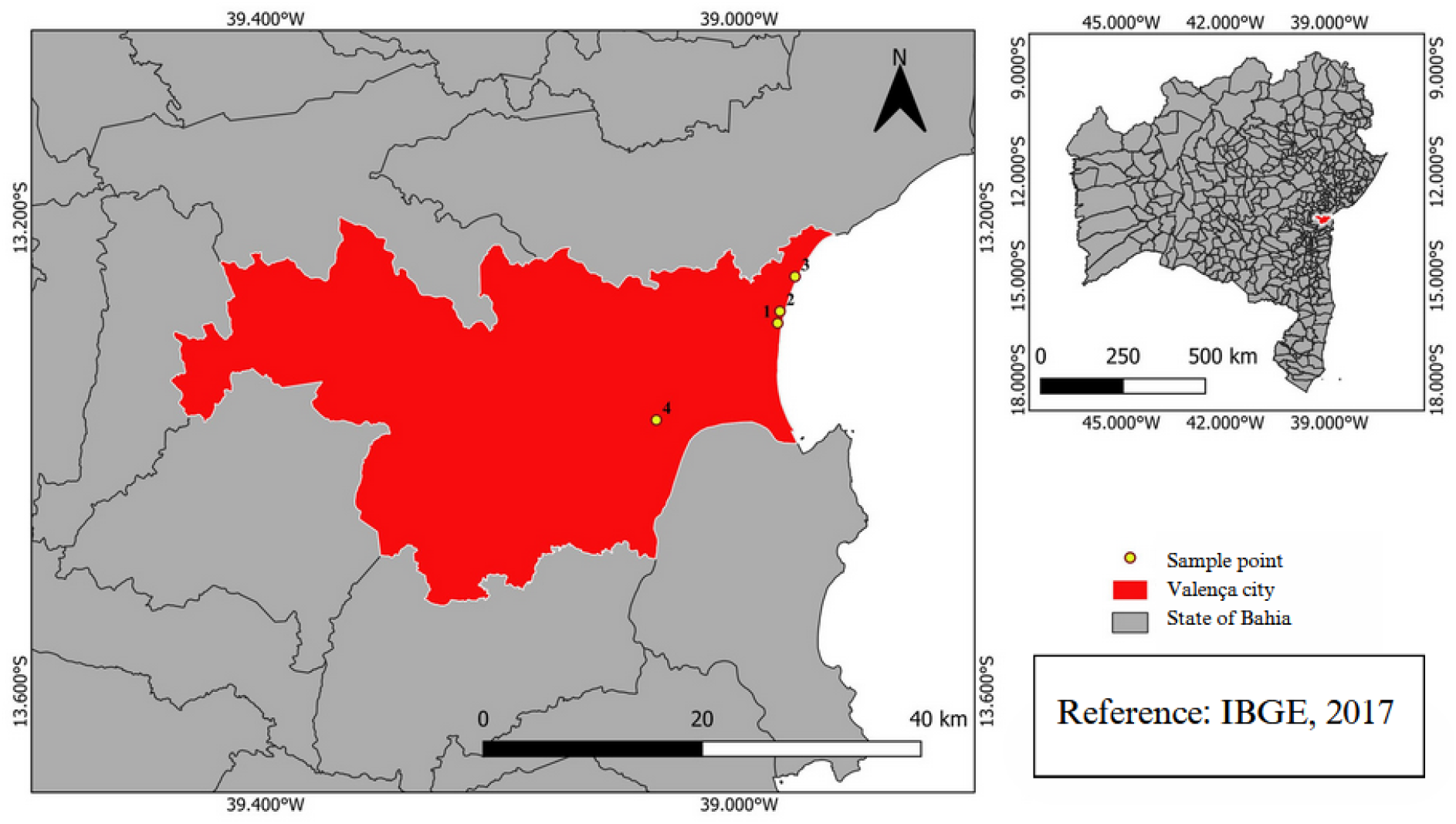
Situation map of Valença city on state of Bahia and sampling points (1. Guaibim; 2. Swamp; 3. Taquari; 4. Una River)

The city has 85.655 citizens and 59,5% of its territory with a suitable sewage system. The predominant sewer system is rudimentary cesspit[25].

Valença has an average altitude of 50 m and a heated and humid tropical weather that favors the action of pedogenic processes. The city developed around the Una River, which forms the local estuarine system. Economic activities are fishing, aquaculture, fluvial transportation for the archipelago of Tinharé, pecuary, oil palm cultivation and tourism, especially in Guaibim district. Many of these activities, in addition to sewage dump, cause various anthropic impacts.

Guaibim is an Environmental Protection Area (APA), mainly formed by mangrove, swamp and restinga ecosystems. Guaibim is located between Jequiriça River and Taperoá channeling, on the coastal zone of Valença.

### Sampling

We did two sampling with a gap of five months, both in low tide, with mild temperature, oscillating between 22 e 24° C. The Figure 2 represents a degradation gradient for the sampling points Guaibim-1 (13°17’27”S, 38°57’54”W), Guaibim-2 (13°16’42”s, 38°57’52”W), Taquari (13°14’59”S, 38°57’13”W) and Una (13°22’14’’S, 39°04’12’’W).

**Figure 2.**
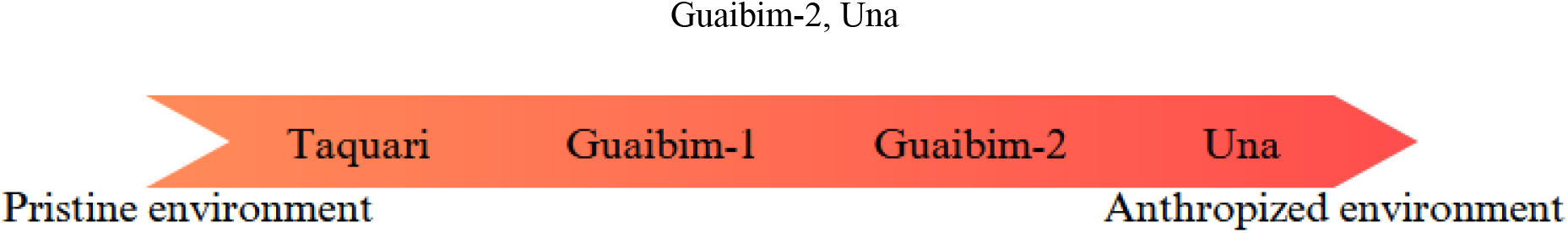
Anthropization gradient of sampling points in crescent order of anthropization: Taquari, Guaibim-1, Guaibim-2, Una

This three first points was collected in Guaibim district and the last one was collected in an urban river in Valença.

In each point, except for Una river, we made a random sampling of seven locals along a transect with 3 m of distance from each other and 10 cm of profundity. We selected these locals to make a comparative between bacterial communities and ARGs presence in pristine and impacted fields.

We homogenized the samples collected in each point in Falcon tubes of 25 ml and involved them in aluminum paper to avoid degradation of DNA and we transported them in polystyrene with dry ice, following recommendations of NeoProspecta, where the metagenomic was realized.

### Extraction of DNA

We extracted the DNA with Quick-DNA/RNA^TM^ MagBead kit (Zymo Research), according manufacturer’s guidelines. We quantified the DNA with a NanoDrop® spectrophotometer (ThermoScientific, USA) and we selected for qPCR compost samples of each point from the first and the second sampling with ratio A260/280 equal or greater to 1.6.

### Genes selection and qPCR

We selected the AMR genes through a list of most used antibiotics on Brazilian healthcare system according literature[26],[27],[28] and informations of Bahia Health’s Secretary. Thus, we selected genes that confer resistance to tetracyclines (*tetA, tetB* e *tetE*), sulfonamides (*sulI*), chloramphenicol (*catA*), quinolones (*qnrA* e *qnrB*) and broad-spectrum beta-lactam antibiotics (*blaSHV* e *blaTEM*). In Table 2 we described the amplified genes, their primers sequence, their size in base pairs and their respective annealing temperature (Ta) in degrees celsius.

**Table 2.**
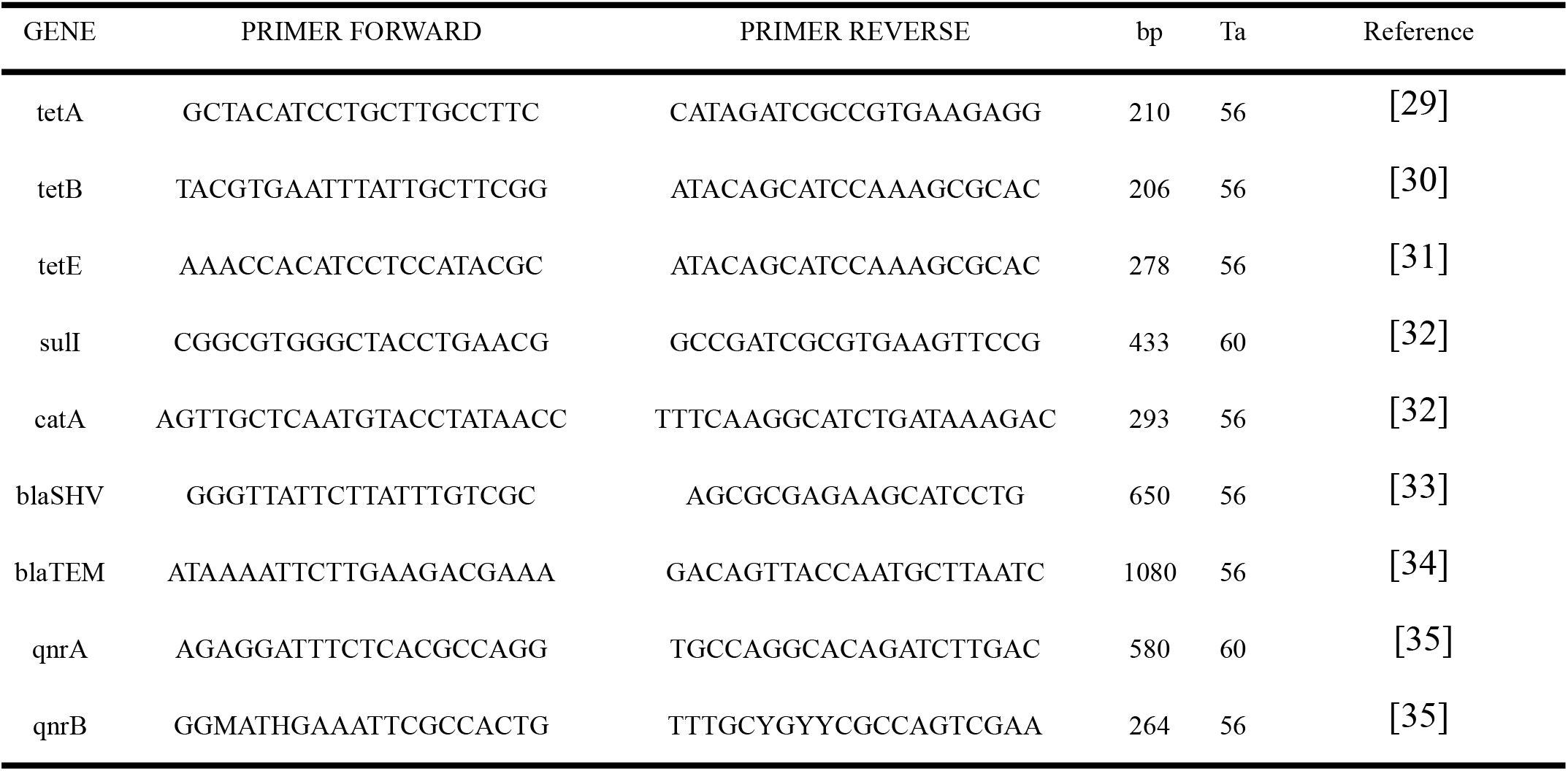
Amplified genes with their respective primers and amplicons (bp).

qPCR of genes considered large, with more than 600 bp is sustained by many authors, as Bassitta et al.[36] e Tolosi et al.[37] Real-time polymerase chain reaction (qPCR) was carried out for the detection of resistance genes. The reagents used and their concentrations for the mix, with a final concentration of 10 μl, are listed in Table 2.

The qPCR mixes, including two negative controls composed of 6 μl of qPCR reagents and 4 μl of nuclease-free water, four quantified standard samples in triplicates, and environmental DNA samples collected at time 1 and time 2, also in triplicates, were distributed into 20 μl microtubes for amplification. For the standard curve construction, we used a serial dilution of synthetic double-stranded DNA (gBlocks®, IDT) from the target 16S rRNA amplicon, with a concentration of 1 ng/μl. The use of this type of fragment as a standard can enhance sensitivity, reliability, and quantification performance[38]. The qPCR was performed on a BIORAD CFX96 Touch System real-time thermocycler, following the programmed steps: 1 cycle at 50°C for 2 min, 95°C for 2 min, followed by 45 cycles at 95°C for 10 s and at the annealing temperature specified in Table 1 for 40 s.

### Molecular identification and metagenomics

Adicionally, we sequenced V3/V4 regions of the gene 16S rRNA, using the primers 341F (CCTACGGGRSGCAGCAG[39] e 806R (GGACTACHVGGGTWTCTAAT[40].

Neoprospecta Microbiome Technologies (Florianópolis, Santa Catarina, Brasil) executed the pipeline.

DNA was extracted using the Quick-DNA/RNATM MagBead kit (Zymo Research, USA) and quantified using the Qubit fluorometer with the dsDNA BR Assay Kit (Invitrogen, USA). Following quantification, the DNA was diluted to 0.5 ng/μL. Genomic libraries were prepared using the V2 kit and sequenced on the MiSeq Sequencing System (Illumina Inc., USA) with 300 cycles and single-end sequencing. The sequences were analyzed using a pipeline employed by Neoprospecta Microbiome Technologies. Sequences with 100% identity were grouped into clusters and used for taxonomic identification by comparison with accurate 16S rRNA sequence databases (NeoRef, Neoprospecta Microbiome Technologies). Sequences with less than 99% similarity to the 16S rRNA gene were excluded from further analyses. The results have been deposited in the GenBank public database (http://www.ncbi.nlm.nih.gov/BLAST/).

### Bioinformatics and data analysis

We transformed the quantification results of ARGs into an order of 10+8, logarithmized, and subjected to R software version 4.2.1 for the construction of bar box plots using the ggplot package.

The reads containing DNA sequences obtained in FASTQ files were loaded into the Galaxy Genome platform (https://usegalaxy.org/) and analyzed using the Mothur pipeline. The quality of the generated sequences was assessed through the FASTQC tool. Subsequently, trimming was performed to remove low-quality reads. The samples were added as replicates, and relative abundance plots and Venn diagrams were generated to study species richness and species co-occurrence in the conducted samplings.

## RESULTS

### Limit of quantification and qPCR efficiency

The quantification limits were established based on the negative control. As the cutoff values for environmental DNA analysis by qPCR are not well-defined in the literature, proprietary methods were adopted to establish these values. In the conducted reactions, the quantification cycle (Cq) values ranged between 22.19 and 42.01, with only three samples exceeding the value of 40. Cq values, even for the investigation of ARGs, above 31 are considered high, especially in clinical settings, and can impact the occurrence of false positives[41],[42]. However, environmental studies have optimized protocols that maintain these higher values for their analyses[37]. Therefore, only reactions amplified up to the 40th quantification cycle were considered for analysis.

We detected resistance genes in various locations at both collection times. The efficiency of the reactions ranged between 93.1% and 100%.

### Absolute concentration and relative abundance of ARGs

All resistance genes were detected in at least one of the sampling locations, but there was variation in both the abundance and frequency of these genes depending on the location. The quantity of each ARGs varied significantly among the sampled locations (p < 0.05).

The quantification results were scaled to a volume of 100 liters and logarithmically (Figure 3-A and 3-B). The DNA concentration for the qnrB, sulI, tetA, and tetE genes was lower, while the qnrA, catA, tetB, blaSHV, and blaTEM genes were found in higher concentrations.

**Figure 3-A:**
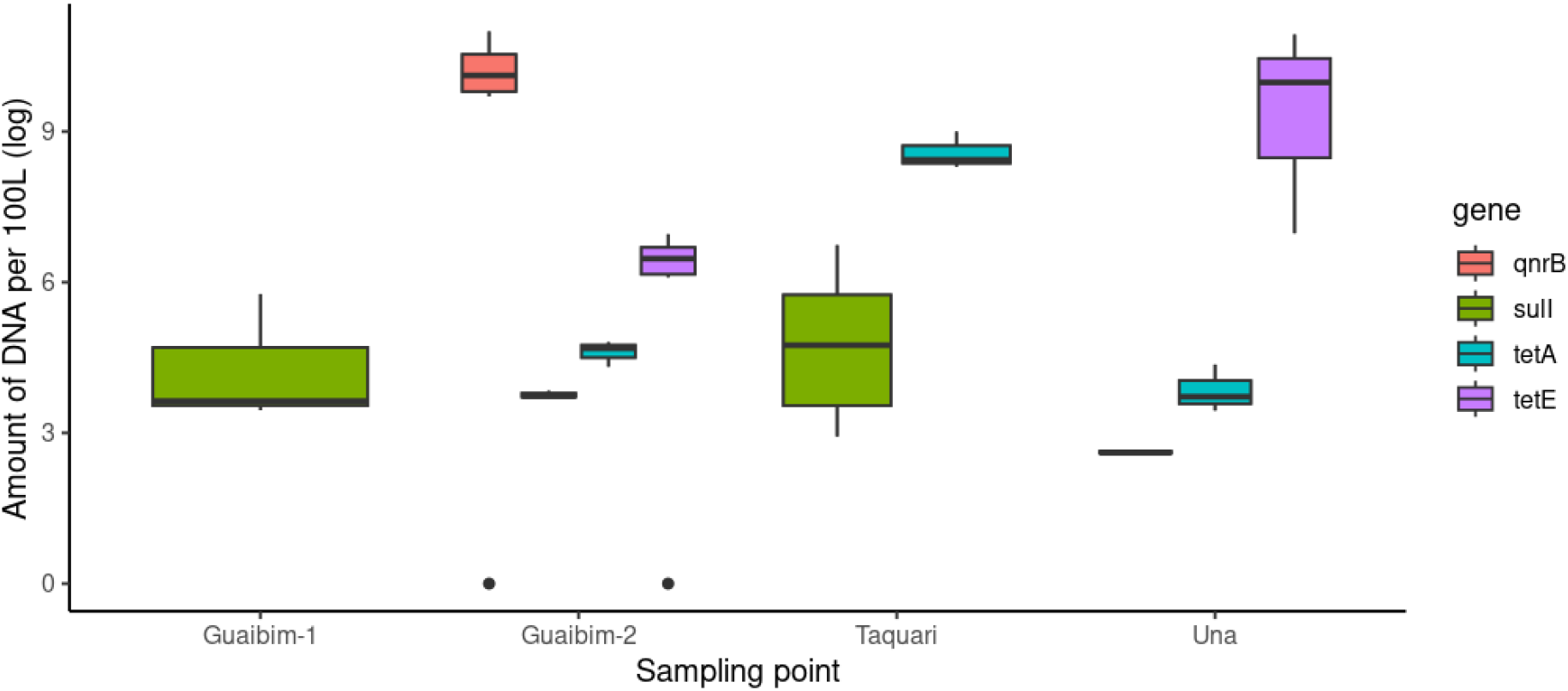
Amount of AGRs (*qnrB, sulI, tetA e tetE*) per 100 litres on both sampling in each sampling point.

**Figure 3-B:**
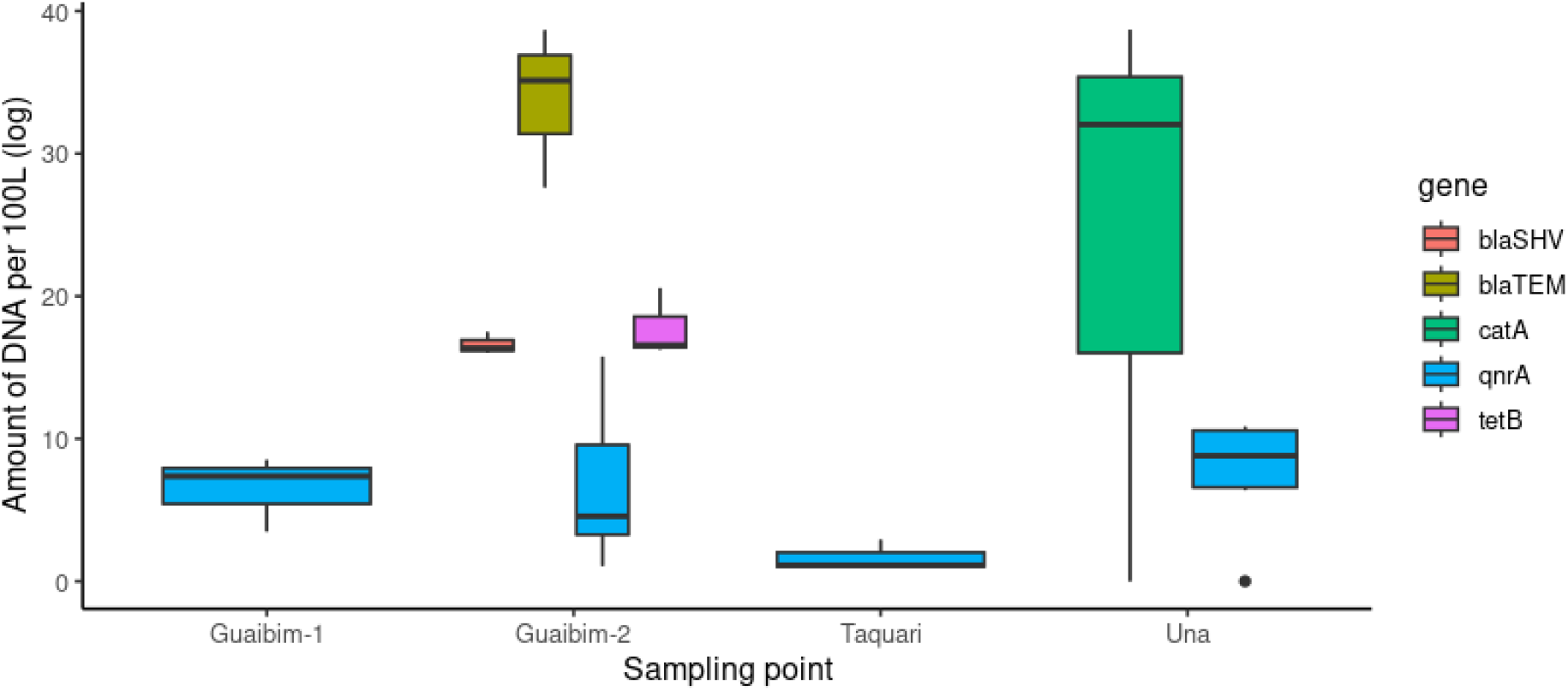
Amount of AGRs (*qnrA, catA, tetB. blaSHV and blaTEM*) per 100 litres on both sampling in each sampling point.

CatA gene was exclusive to samples collected from the Una River, with significant variation observed among the replicates. On the other hand, the qnrB gene was exclusive to samples collected in Guaibim-2, where it can also be noted that the variation in DNA quantity among replicates was lower.

Figure 4 illustrates the relative abundance of ARGs on the four collection points. Samples collected in Guaibim-2 exhibited a higher quantity of ARGs. Guaibim and Taquari showed the lowest quantities, with limited diversity of ARGs. In the samples collected from Una, an intermediate quantity was observed compared to the aforementioned points, with almost half of this quantity attributed to catA. TetA was detected in all samples, and the sulI gene also showed a widespread distribution.

**Figure 4:**
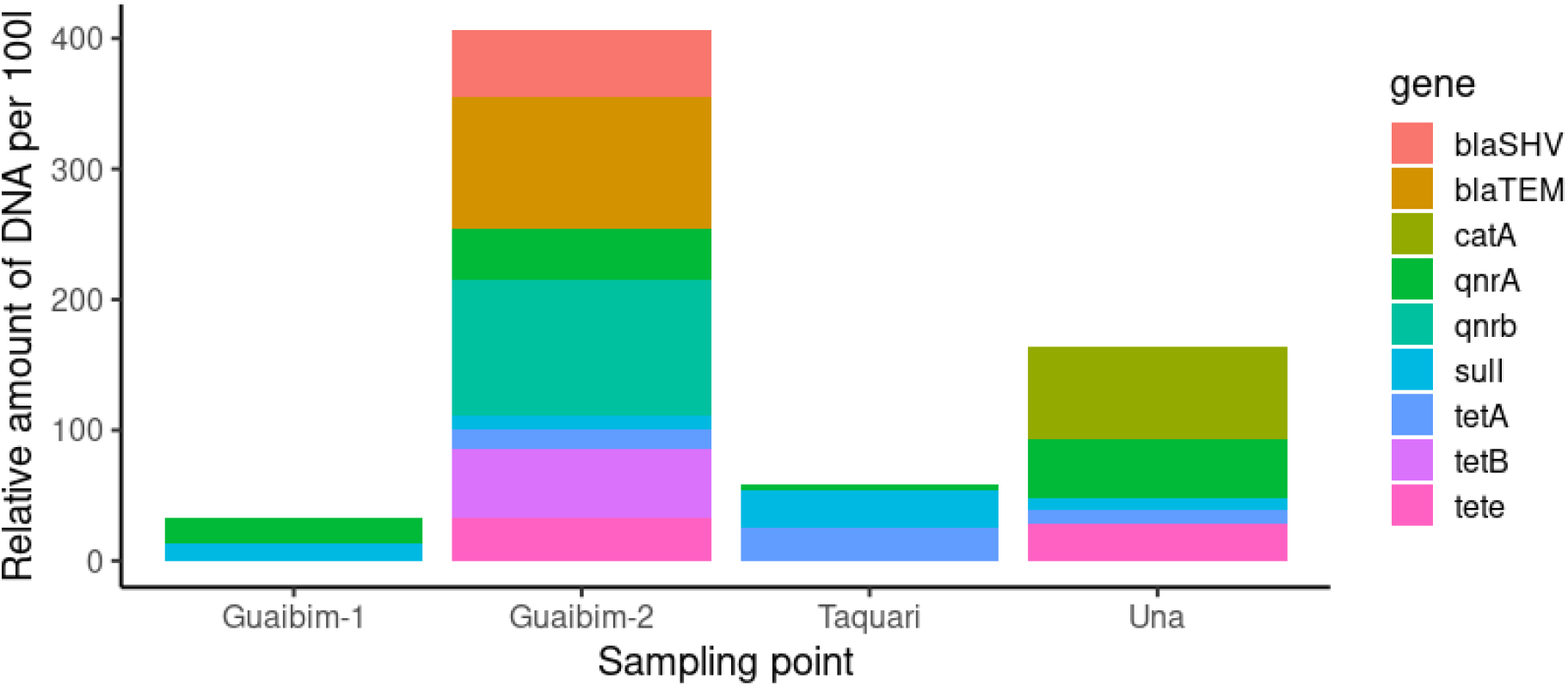
Relative amount of ARGs per 100 L in each sampling point.

### Bacterial community composition

We identified a total of 60,000, 36,755, 37,746, and 54,248 sequences in the samples collected in Guaibim-1, Guaibim-2, Taquari, and Una, respectively. The majority belonged to the phylum Proteobacteria (51%), followed by Firmicutes (8%), Bacteroidetes (8%), Actinobacteria (5%), Chloroflexi (4%), and Acidobacteria (4%). About 17% of the bacteria could not be classified (Figure 5). Within Proteobacteria, notable groups include Gammaproteobacteria, Alphaproteobacteria, and Deltaproteobacteria.

**Figure 5:**
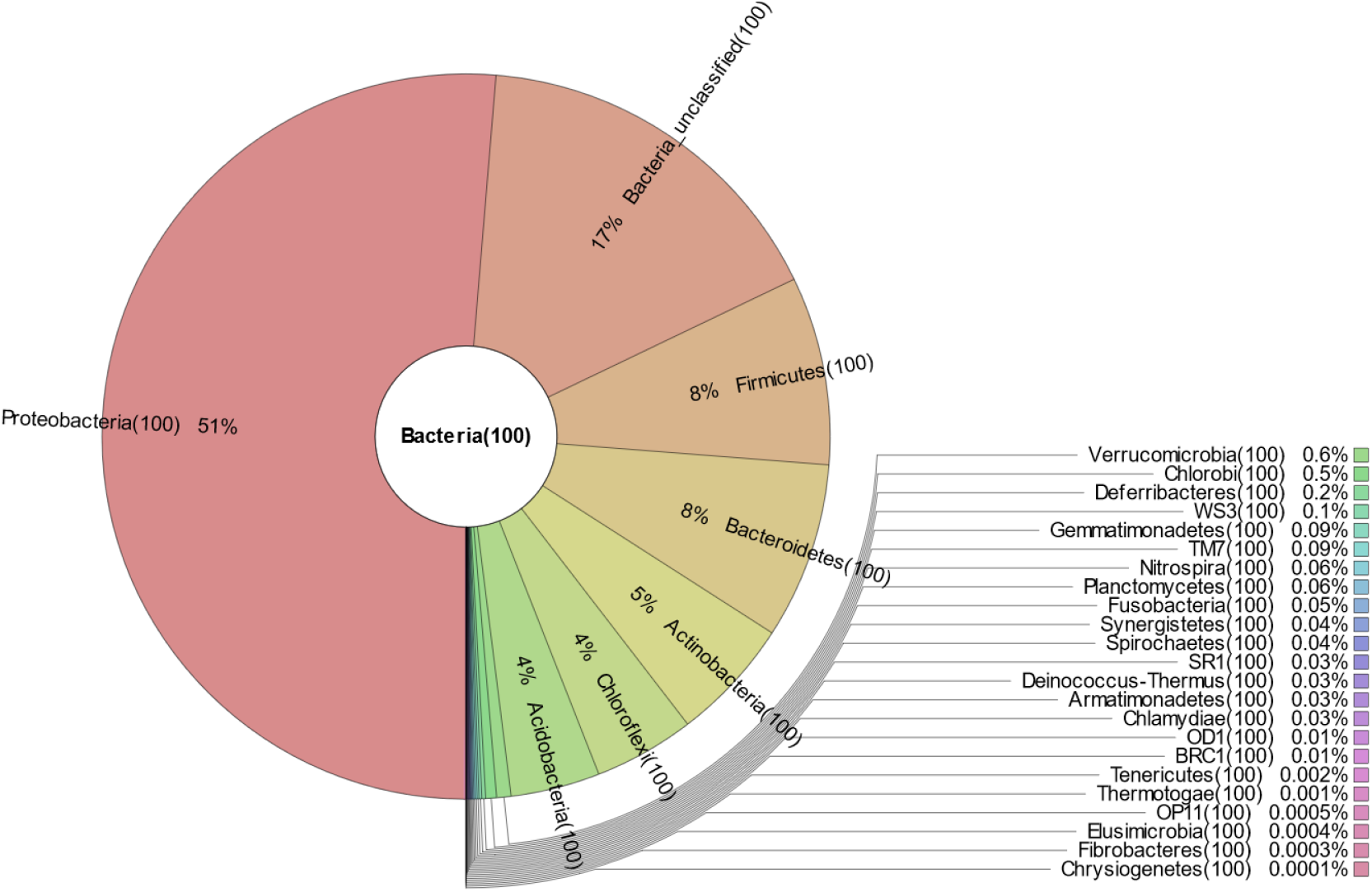
Relative frequency of bacterial groups in the sampled communities.

The relative frequency of these groups varied at each collection point, but the main bacterial groups were represented in all samples (Table 3).

**Table 3.** qPCR reagents.

| Reagent | µl per reaction |
| --- | --- |
| PowerUp™ SYBR® Green Master Mix (Thermo Fisher Scientific, USA) | 5 |
| Primer forward (100pmol) | 0.5 |
| Primer reverse (100pmol) | 0.5 |
| DNA (10ng) | 4 |

**Table 3.** Abundance of the main phyla comprising the microbial communities at each sampling point (expressed in the number of sequences).

| Phyla | Sampling Point |  |  |  |
| --- | --- | --- | --- | --- |
|  | Guaibim-1 | Guaibim-2 | Taquari | Una |
| Proteobacteria | 44819 | 17377 | 26427 | 24121 |
| Actinobacteria | 4811 | 2779 | 2891 | 4971 |
| Firmicutes | 1482 | 7938 | 523 | 6229 |
| Chloroflexi | 2757 | 891 | 2388 | 5924 |
| Bacteroidetes | 1640 | 1315 | 1327 | 6911 |
| Acidobacteria | 486 | 2015 | 452 | 357 |
| Não identificadas | 3948 | 2165 | 2694 | 7627 |

Additionally, we observed that samples collected from Una exhibited a higher number of exclusive species. Guaibim-1 and Taquari showed a higher co-occurrence of species, followed by Guaibim-2 and Una (Figure 6)

**Figure 6:**
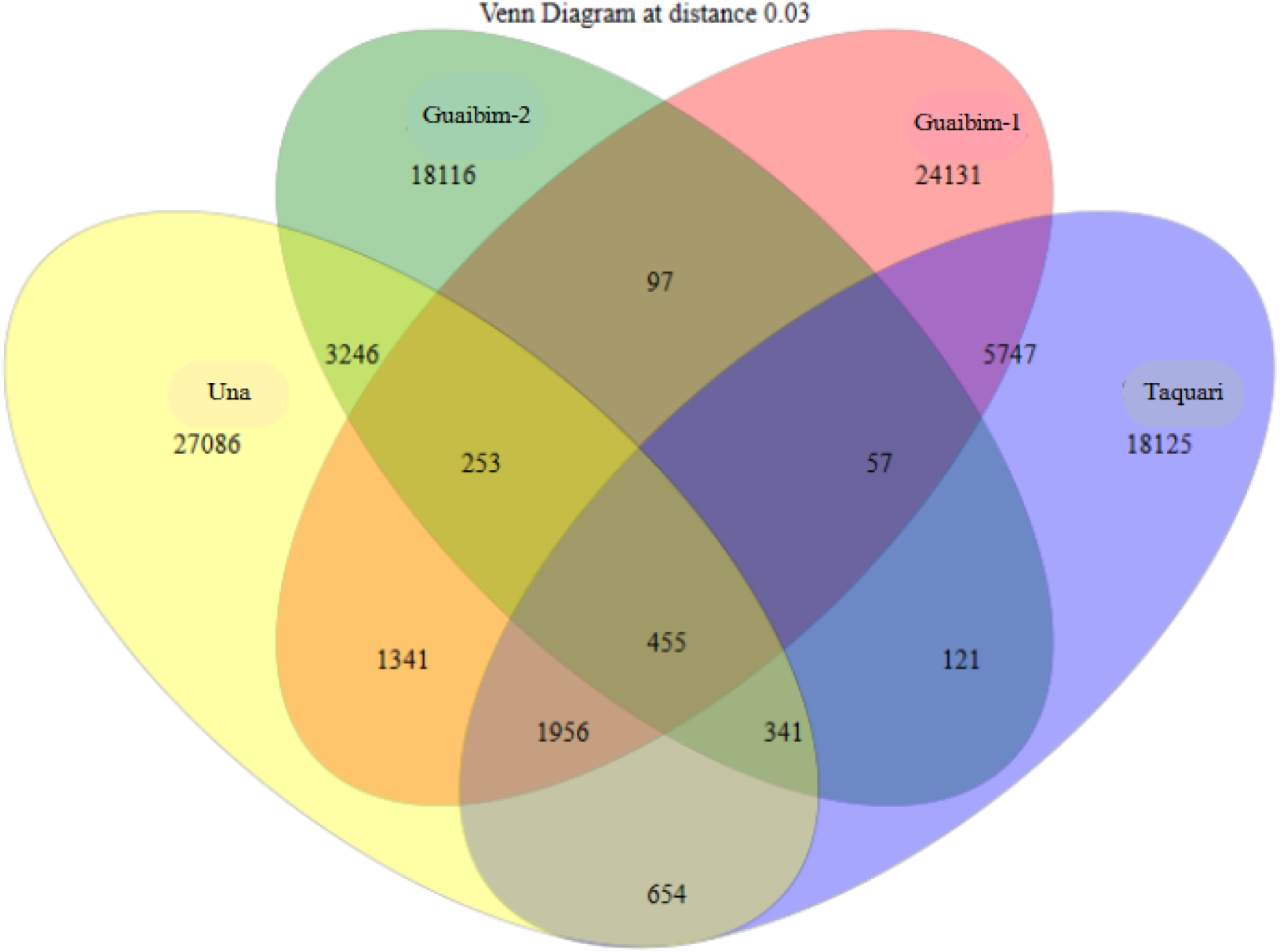
Venn diagram demonstrating co-occurrence on sampling points

### Presence of pathogenic bacteria

Initially, we verified the presence of bacteria from the ESKAPE group (*Enterococcus faecium, Staphylococcus aureus, Klebsiella pneumoniae, Acinetobacter baumannii, Pseudomonas aeruginosa*, and *Enterobacter spp*). These bacteria are listed by the World Health Organization as a priority for surveillance. They are often associated with multidrug resistance, have rapid replication, high transmissibility, and frequently cause prolonged infections due to the difficulty of treatment[43].

In Guaibim-1, we detected sequences belonging to the groups *Enterococcus faecium* and *Acinetobacter* sp. In the samples collected in Una, *Klebsiella pneumoniae, Enterococcus faecium*, and *Enterobacter cloacae* were found. Other bacteria causing infections were also identified, such as *Escherichia coli*, common in the intestinal tract of various animals, although associated with infections in the urinary system, and *Clostridium botulinum*, the bacterium that causes botulism.

In addition to pathogenic bacteria, we observed a high frequency of bacteria associated with sulfur cycling and methanogenic bacteria.

## DISCUSSION

The diversity of ARGs in Guaibim-2 may be related to the lack of an adequate sanitation system in the region. As waste is directed to septic tanks, it is possible that these genes are released by local residents, and the overflow from septic tanks may contaminate the waters that make up the swamp ecosystem. Additionally, the use of water by animals to which antibiotics are administered, creating a habitat with significant pressure for the selection of resistant bacteria, may be another contributing factor to the increased diversity of ARGs at this sampling point.

In Taquari, Una, and Guaibim-1, there is also the influence of tides. Although estuaries are deposition environments, tidal variation may be a factor contributing to a greater dispersion of resistance genes, as observed in other studies[44],[45], while Guaibim-2 constitutes a more stable environment, with a higher rate of deposition of ARGs.

Microbial community composition, with a higher frequency of bacteria from the phyla Proteobacteria, Actinobacteria, and Firmicutes, is similar to previously published studies[46,47]. In general, there is a wide diversity of deltaproteobacteria, including families such as *Desulfobacteraceae* and *Desulfobulbaceae*, which participate in the reduction of sulfates to sulfites, and others involved in methane metabolism. Various groups belonging to the phyla Alphaproteobacteria, Betaproteobacteria, and Deltaproteobacteria, which are abundant in all samples, play important roles in nitrogen cycling[13].

Previously published research indicates a higher representation of Bacteroidetes in soils contaminated with petroleum.[48]. However, the frequency of Bacteroidetes was low in all samples. There is also a correlation between the presence of anaerobic bacteria and environmental contamination[49]. From this perspective, it is evident that the higher quantity of bacteria from methanogenic families such as *Methanomicrobiaceae, Methanosarcinaceae, Methanococcaceae, and Methanobacteriaceae*, as well as *Desulfovibrionaceae* and *Desulfobacteraceae* in samples collected from Guaibim-2 and Una, indicates a higher level of contamination in these locations. It is also important to consider the impact of greenhouse gas production on regional climate concerns.

Additionally, bacteria from the phylum Firmicutes are often associated with the degradation of polycyclic aromatic hydrocarbons. Their high frequency in Guaibim-2 and Una may indicate the presence of these compounds, which positively influenced the selection of organisms capable of metabolizing them. The diverse metabolic pathways of Firmicutes make them a group widely used in bioremediation processes for areas contaminated with hydrocarbons, industrial waste, and as pest controllers[50,51].

The higher proportion of bacteria belonging to the phylum Acidobacteria is indicative of soil pH variation. Although naturally occurring and widely distributed, the relative frequency of this group in microbial communities is low, tending to increase in more acidic soils[52]. Pedrinho et al. demonstrated that the use of sewage sludge tends to increase the frequency of Acidobacteria and reduce community diversity[52]. “In light of this, it is crucial to note that, while the relative abundance of Acidobacteria is three times higher in the swamp compared to other locations, the total DNA quantity from this group is not significant enough to suggest a high sewage contamination rate in this region. Nevertheless, when coupled with the observation that the diversity of ARGs detected at this sampling point is lower, it underscores the importance of ongoing site monitoring.

In general, regarding the effects of contamination on the microbial community composition at the phylum level, little can be inferred from metagenomic analyses for this study. Various studies indicate that different environments may have distinct biotic compositions, which vary according to various physicochemical parameters (pH, available nutrients, salinity, and various types of pollutants). For a more accurate assessment, continuous monitoring considering these parameters would be necessary, examining changes in the microbiota composition at the sampled locations after different events.

Metagenomics, in this context, serves as a crucial source of supplementary data for investigating antimicrobial resistance in a given location. Indeed, some bacteria listed as a priority for surveillance by the World Health Organization were detected in the samples. Enterococcus faecium, Acinetobacter sp, Klebsiella pneumoniae, Enterococcus faecium, and Enterobacter cloacae have a documented history of resistance against potent antibiotics commonly employed in healthcare systems[22,52]. The co-occurrence of these bacteria with ARGs, especially their presence in locations where a higher diversity of resistance genes was found, indicates that Guaibim-2 and Una are areas of public health concern. This is particularly noteworthy as these locations may serve as sources of water and food resources for humans or animals.

The substantial overlap in species between Guaibim-1 and Taquari suggests the close proximity of these environments, indicating a higher degree of microbial community similarity. Conversely, Guaibim-2 and Una exhibit distinct characteristics and are not geographically adjacent. This leads to the inference that their preservation status, likely influenced by the presence of domestic effluents and hydrocarbons (such as microplastics), has been a contributing factor in selecting for the observed bacterial groups.

Furthermore, both Guaibim-1 and Taquari had limited diversity of resistance genes, suggesting that neither of them has reached highly concerning contamination levels. The absence of bacteria from the ESKAPE group (as well as other groups of infectious organisms commonly found in sewage) can also be considered an indicator of the low contamination level.

Considering the abundance of DNA from these genes in Guaibim-2 and Una, especially the likelihood of natural origin for the observed resistance is minimal. Notably, the antibiotics associated with the ARGs identified in this study are commonly used in the public health network of Bahia, suggesting potential effects on the selection of ARGs in various studied locations, even those presumed to be preserved environments. However, due to insufficient data in this study to trace the origin of antimicrobial resistance, it is not possible to conclusively attribute the high frequency of detected ARGs to the use of antibiotics in livestock, aquaculture, or human health. It is crucial to highlight that, even in areas lacking ESKAPE group bacteria, the elevated frequency of ARGs and the presence of other infection-causing bacteria indicate that untreated water is unsuitable for human consumption, posing a risk for the transfer of resistance genes[53].

## CONCLUSIONS

The widespread distribution of ARGs indicates high antibiotic usage for the treatment of bacterial infections in the Valença region, with tetracyclines and quinolones being the predominant classes. The data highlight the urgency for the development of integrated studies involving environmental and health agencies to monitor microbial resistance. Additionally, it underscores the need for awareness campaigns regarding the consumption of untreated water and the indiscriminate use of antibiotics.

The presence of bacteria indicating contamination from human waste in the environment is alarming and signals deficiencies in the sanitation sector in the city, especially in Guaibim, where a sanitary sewer system has not been installed yet. It is noteworthy that, even in Valença, where a sewage system exists, albeit not fully, ARGs were detected at the sampling point. This underscores the need for the development of tools and techniques capable of treating and eliminating resistance genes from waste discharged into water bodies and their integration into the sanitation system.

Furthermore, there is a need for studies to assess the presence of ARGs in the region’s fisheries, which are frequently consumed and traded. Through continuous monitoring of these genes in food, coupled with effective treatment of domestic and industrial waste and the judicious administration of antimicrobial medications in healthcare systems, it is possible to break the cycle of gene transmission and mitigate future outbreaks of infections caused by resistant organisms.”

## ACKNOWLEDGEMENTS

Thanks to the State University of Feira de Santana, the Graduate Program in Ecology and Evolution, to the Bahia State Research Support Foundation (FAPESB) and Coordination for the Improvement of Higher Education Personnel (CAPES).

